# Genealogical inference and more flexible sequence clustering using iterative PopPUNK

**DOI:** 10.1101/2022.10.08.511450

**Authors:** Bin Zhao, John A. Lees, Hongjin Wu, Chao Yang, Daniel Falush

## Abstract

Bacterial genome data are accumulating at an unprecedented speed due the routine use of sequencing in clinical diagnoses, public health surveillance and population genetics studies. Genealogical reconstruction is fundamental to many of these uses, however, inferring genealogy from large-scale genome datasets quickly, accurately, and flexibly is still a challenge. Here, we extend an alignment- and annotation-free method, PopPUNK, to increase its flexibility and interpretability across datasets. Our method, iterative-PopPUNK, rapidly produces multiple consistent cluster assignments across a range of sequence identities. By constructing a partially resolved genealogical tree with respect to these clusters, users can select a resolution most appropriate for their needs. We demonstrated the accuracy of clusters at all levels of similarity and genealogical inference of iterative-PopPUNK based on simulated data and obtained phylogenetically-concordant results in real datasets from seven bacterial species. Using two example sets of *Escherichia/Shigella* genomes and *Vibrio parahaemolyticus* genomes we show that iterative-PopPUNK can achieve cluster resolutions ranging from phylogroup down to sequence typing (ST). The iterative-PopPUNK algorithm is implemented in the ‘PopPUNK_iterate’ program, available as part of PopPUNK package.

## Introduction

A key step in many analyses of collections of bacterial genome sequences is to understand how the sequences are related to each other. For example, to make a clinical diagnosis, a key question is which previously characterized organisms are similar to strains present in the patient sample (Gardy and Loman 2018; Gu et al. 2019). To identify transmission chains, it is necessary to establish which of the sequenced strains share very recent common ancestors, indicating a shared source (Croucher and Didelot 2015; Sintchenko and Holmes 2015). To map genotype-phenotype relationships, it is often helpful to delineate lineages. Organisms from the same lineage have similar sequence content and thus are likely to have similar phenotypes (Falush 2016; Power et al. 2017; Brinda et al. 2020). To accurately reconstruct the evolutionary history of a species, it is necessary to obtain a sample of organisms that is broadly representative of the species’ diversity as a whole and is not biased towards specific lineages, for example by containing only multiple closely related sequences from the same environmental source.

A “gold standard” for establishing genetic relationships between strains is through genealogical reconstruction. Any given sample of bacteria are related to each other by a tree, with each node of the tree representing the most recent common ancestor (MRCA) of the strains below it in the genealogy. The genealogical history reflects the clonal relationships between strains; it is unique and unambiguous. Genome sequence data provides partial information on the topology of this tree, its MRCA and where and when that MRCA existed. In practice, the information that genome sequences provide on genealogical relationships is both incomplete and time consuming to extract. Moreover, a full genealogy is rarely necessary for most practical uses of genetic relationships. This has led to the development of methods that attempt to identify clusters of closely related organisms. A simple interpretation of each cluster is that it corresponds to the set of strains descended from a single node of the genealogical tree. Amongst these possible methods, PopPUNK is notable because it works from raw sequence data and does not require a genome alignment (Lees et al. 2019). This means that the clustering can be obtained with significantly less computational resources and is much faster than methods based on sequence alignments. Moreover, PopPUNK is easily extendable, as new batches of genomes can be integrated into an existing database without needing to recalculate all pairwise distances and reperform the model fitting step.

PopPUNK avoids the need for an alignment by creating a *k*-mer profile for each strain, using both short and long *k*-mer lengths, which is computationally much faster than any form of alignment thanks to highly efficient hashing algorithms. It compares the *k*-mer profiles of different strains based on their Jaccard similarity. Gene presence-absence polymorphisms create a profile of *k*-mer similarities which is distinct from that due to SNPs, making it possible to estimate distances for core and accessory genome separately by fitting the pattern of distance profiles across multiple *k*-mer lengths.

Two machine learning algorithms, Bayesian Gaussian Mixture Model (BGMM) and Hierarchical Density-Based Spatial Clustering of Applications with Noise (HDSCAN), are implemented in PopPUNK to identify components. The component closest to the origin of the scatterplot of core and accessory distances is defined as “within strain”, corresponding to closely related strains. The other one is defined as “between strain”, corresponding to distant strains. Then these within-strain distances are used as edges to link samples and create PopPUNK clusters. The PopPUNK algorithm identifies a single level of clusters, attempting to find an “optimal” choice. However, the biological interpretation of this optimal clustering is not clear since the genealogical tree underlying the relationships between strains is hierarchical rather than having a single level. Furthermore, in practice, multiple levels of clusters with different genetic similarities are often required to achieve different purposes. For example, in the disease outbreak scenario, epidemiologists require clusters of very closely related or indistinguishable strains to help identify and track outbreaks. For routine surveillance and subtyping, clusters with higher diversity are required to represent strains from a sequence type or clonal group. Population geneticists usually require clusters of population or sub-population level to conduct species-level studies.

Two state-of-the-art pipelines for large-scale genome datasets that allow multi-level clustering are Fastbaps (Tonkin-Hill et al. 2019) and HierCC, as implemented in EnteroBase (Zhou et al. 2021). Fastbaps is based on a core genome alignment, which is slow to generate for datasets of even moderate sizes. HierCC requires an appropriate curated database of genes and alleles or a stable strain nomenclature and cannot be the general solution for most bacterial species.

Here we present iterative-PopPUNK – an extension of the PopPUNK software - that allows estimation of a partially resolved genealogical tree and provides clustering consistent with that tree at different levels of sequence similarity. This enhancement provides greater flexibility than the original PopPUNK algorithm, allowing the fineness of the clustering to be adjusted according to the purpose at hand. Moreover, the clusters can be interpreted more readily, by reference to the full (partially resolved) genealogy.

## Results

### Overview of iterative-PopPUNK

Iterative-PopPUNK allows estimation of a partially resolved genealogical tree and provides clustering consistent with that tree at different levels of sequence similarity. This enhancement provides greater flexibility than the original PopPUNK algorithm, allowing the fineness of the clustering to be adjusted according to the purpose at hand. Moreover, the clusters can be interpreted more readily, by reference to the full (partially resolved) genealogy. The speed of iterative-PopPUNK is similar to PopPUNK, which can complete the analysis of thousands of bacterial genomes in a few minutes, and can scale even to very large-scale genome datasets (e.g. >100,000). Using simulated data, we demonstrate the accuracy of the clusters from iterative-PopPUNK at all output levels of similarity. Furthermore, we show that for real data, the produced clusters are normally highly concordant with those obtained by phylogenetic methods, but without requiring an alignment.

An overview of Iterative-PopPUNK pipeline is shown in Figure 1A. Six steps are required to implement the algorithm of iterative-PopPUNK. The first two steps proceed as usual using the original PopPUNK algorithm: 1) database construction and distance calculation and 2) model fitting using original PopPUNK algorithm. Iterative-PopPUNK then performs four additional steps to achieve multi-level clustering and generate clusters under the given similarity cutoffs.

**Figure 1:**
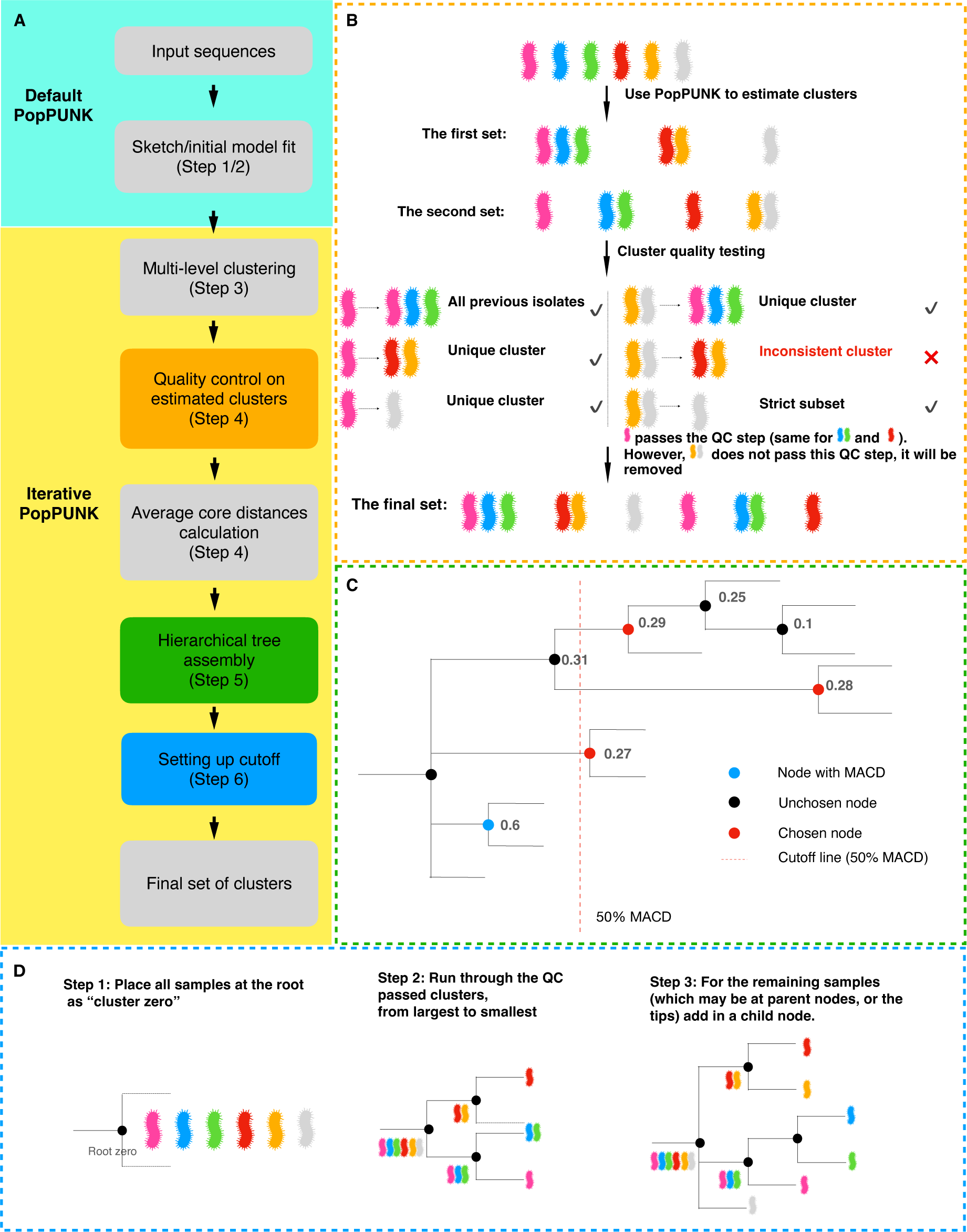
Workflow for developing iterative-PopPUNK. A). Steps for designing iterative-PopPUNK. After inputing sequences data, PopPUNK creates local sketching database which is furthur used to estimate clusters. QC steps for removing inconsistent clusters is shown in orange box. Green box shows how to nest these estimated clusters into an iterative-PopPUNK tree. Methods for choosing final set of clusters are presented in blue box; B). QC algorithm. Three conditions for determing QC-passed clusters are discribed in details in methods, which in short are: 1) the new clusters contain all of isolates from previous clusters; 2). the new cluster is unique and contains none of items from previous clusters; 3). the new cluster is a strict subset of one of previous clusters. C). Algorithm for cutting iterative-PopPUNK tree. The goal is to choose the closest node to the cutoff line but with a smaller value. In this example, the cluster (node) annotated using blue color has the maximum value of average core distances (0.6). The red dashed line shows the cutoff is 50% of the MACD (0.3). Therefore, the node with an ACD lower than and closest to 0.3 will be selected. D). Hierachical tree assembly. The dashed lines indicated these potential branches during the tree assembly process.

### Step 3: Multi-level clustering by moving decision boundary iteratively

After an initial model has been fitted, PopPUNK identifies two fitted components representing “within-strain” comparisons between closely related isolates and “between-strain”. However, unlike the original PopPUNK algorithm, which identifies a single boundary for defining a “best” cluster assignment by optimising a heuristic (the network score), iterative-PopPUNK instead ignores the network score, and places the decision boundary at multiple equally spaced positions perpendicular to the line connecting within- and between-strain components to create multi-level cluster assignments. We have found using simulated data that the clustering accuracy remains high even when the network score is low and we are interested in obtaining clustering at multiple hierarchical levels.

The first viable boundary point is where at least one non-trivial cluster (containing at least two samples) is formed, corresponding to the two most closely related sequences in the dataset. The number of boundaries, and therefore number of starting sets of clusters can be set by the user. Based on simulation data, we found that the number of estimated clusters increases as the number of decision boundary line increases, and basically reached a plateau when the number of decision boundary line is 30 (Supplemental Fig S1). To balance the speed and resolution of iterative-PopPUNK, we set 30 as the default cutoff. Users can choose to increase this value if they want a higher resolution.

### Step 4. Quality control (QC) of clusters obtained at different levels

QC is performed to remove any duplicated or inconsistent clusters. The clusters obtained under the first viable boundary point are directly defined as QC-passed clusters due to their high accuracy based on the evaluation using simulation data. A new cluster assignment C_j_ (which is a set of samples) at boundary point *j* was compared with QC-passed clusters C_i_ at the previous boundary *i*, and is considered QC-passed if and only if it meets one of the following conditions, as shown in the conceptual outline in Figure 1B:

1) C_j_ contains all of isolates from a QC-passed cluster C_i_ (C_i_ ⊆ C_j_, condition 1);
2) C_j_ is unique and contains none of isolates from any previous C_i_ (Cj ν Ci = 0, condition 2);
3) C_j_ is a strict subset of one of the previous QC-passed cluster C_i_ (C_j_ ⊆C_i_, condition 3);

By iterating through all pairs of clusters between the current step and next step forward (as clusters increase in size; equivalently the boundary moves upwards), a new QC-passed C_j_ was added into the list C_i_ at the next round, until the QC of all the clusters is completed.

### Step 5. Partially resolved genealogical tree construction and average core distance calculation

A partially resolved genealogical tree is then constructed based on QC-passed clusters by nesting clusters within larger ones (Figure 1D). The algorithm we used to do this is as follows:

1) Place all strains at the root as “cluster zero” and sort all clusters in a descending order according to their size;
2) Run through the QC-passed clusters, from largest to smallest. For each one:

a) Look through all clusters which have been added to the tree and find the smallest cluster which is a superset of the cluster being tested;
b) Nest the new cluster as a child under this parent cluster;
c) Remove the strains in the child cluster from the parent cluster;
3) For the remaining strains (which may be at parent nodes, or the tips) add in a child node;
4) For each of the nodes in the tree, we calculate the average core distances between the strains under this node and set its branch length relative to root accordingly.

### Step 6. Choosing clusters under given similarity cutoffs

Considering the large difference in the genetic diversity of different bacterial species, we designed two types of relative cutoffs to use when quantifying genetic similarity within a species: 1) P_max-cluster-dist_, the percentage of Maximum value of the Average Core-Distance (MACD) of all the iterative-PopPUNK clusters (extracted from local PopPUNK distance database). The cluster with maximum value of the MACD has the highest genomic diversity. We use the percentage of this value as a cutoff to decide which other clusters are selected. 2) P_bet-mean-dist_, the percentage of mean pairwise core-distance between all isolates (extracted from initial model fitting results);

When one of these cutoffs is provided, iterative-PopPUNK searches all the nodes except root node in the partially resolved tree and extracts the relative similarity values (P_max-cluster-dist_ or P_bet-mean-dist_) of all the samples under each node (NP) and its parent node (N+1P). If a node meets the condition NP ≤ cutoff < N+1P, all the child strains under this node are assigned into a cluster (Figure 1C). The remaining unassigned strains were defined as non-clustered strains.

### Evaluation of iterative-PopPUNK using simulation data

To test the performance of iterative-PopPUNK, we generated seven datasets of simulated genomes where the true genealogy is known. We generated 100 genomes with a length of 1,000 kb per dataset and their corresponding genealogical trees using FastSimBac (De Maio and Wilson 2017), representing bacteria with different mutation and recombination rates (μ = 0.01-0.06, r=0.003-0.04, Supplemental Table S1). We created clusters of these simulated genomes using PopPUNK and iterative-PopPUNK using three different types of model fit (HDBSCAN and BGMM with two and three components), and mapped the clusters obtained back to the simulated genealogical trees.

To assess performance, we compared the consistency of the output clusters with the true simulated clonal relationships, which are represented in genealogical trees (Figure 2A and 2B). Nodes of the tree that correspond exactly to an inferred cluster are shown in red. Where an inferred cluster does not correspond to a node in the true tree, it is assigned a colour and all the strains in that cluster have their tips labelled using that colour. For example, for simulated data obtained with a recombination rate (r=0.001) and mutation rate (μ=0.001, Figure 2A, 2B), PopPUNK and iterative-PopPUNK (after step 4) generated 9 and 69 clusters, with the accuracy of 100% (9/9) and 99% (68/69), respectively. This criterion is stringent but strains from unmatched clusters were closely related in the simulated tree (Figure 2B).

**Figure 2:**
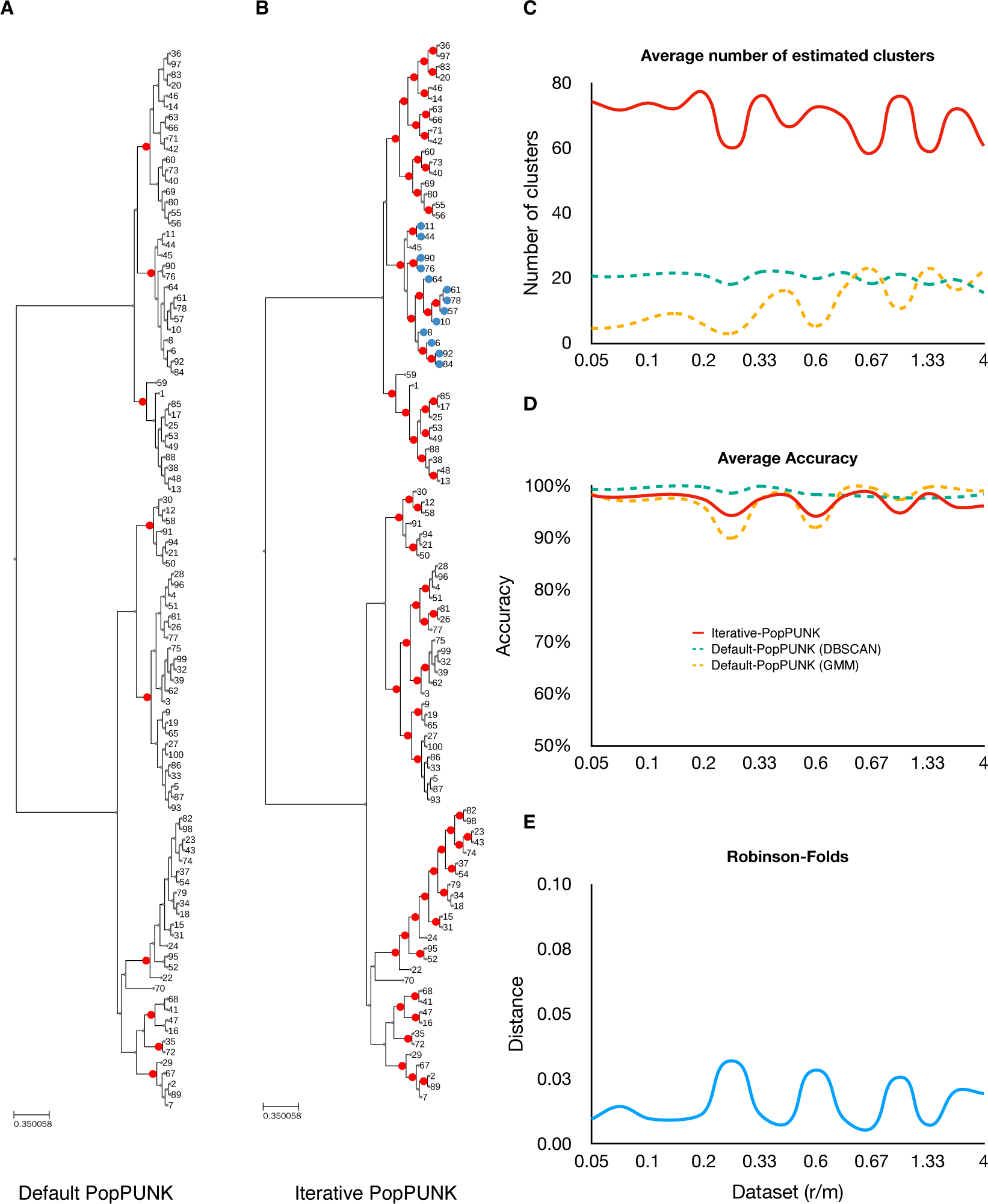
The performance of iterative-PopPUNK on simulated data. The chosen simulated dataset for A and B has a recombination rate r=0.001 and mutation rate μ=0.01. A). A genealogical tree with 9 nodes (clusters) estimated by default PopPUNK; B). The same genealogical tree with 69 clusters estimated by iterative-PopPUNK, among which 68 can be totally matched with genealogical tree nodes while 1 of them cannot be matched, with strains in the unmatched cluster shown in blue; C). A line graph shows the average number of estimated clusters by PopPUNK and by iterative-PopPUNK. The solid lines present the results for iterative-PopPUNK, and the dash lines are for default-PopPUNK. D). The accuracy results for both default PopPUNK and iterative-PopPUNK. The accuracy is calculated from total number of matched nodes divided by total number of estimated clusters. E). Robinson Folds distances between true trees and iterative-PopPUNK trees. r/m stands for recombination rate is divided by mutation rate.

Overall, iterative-PopPUNK generates many more (on average ∼6-fold greater) clusters than PopPUNK with similar accuracy across a range of different mutation and recombination rates (Figure 2C and 2D). A universal model we can recommend for iterative-PopPUNK is BGMM with two components (K = 2), because more datasets could be analysed using this setting (HDBSCAN fits sometimes fail to converge). Under this model, the average accuracy of clusters remains high across parameter choices (∼95%). The average Robinson Folds distance between true trees and iterative-PopPUNK trees is very low (∼0.02), which is in highly concordance with our own accuracy testing method (Figure 2E). Therefore, we set it as the default model of iterative-PopPUNK and applied it in further analysis.

We next tested the performance of iterative-PopPUNK under different relative genetic similarity (P_max-cluster-dist_) cutoffs. We again use the results obtained for simulated data under a mutation and recombination rate (μ=0.001, r=0.001, Figure 3A) as an example, and mapped iterative-PopPUNK clusters under different cutoffs back to simulated trees. As expected, the genetic diversity of strains within a cluster increases as the cutoff increases. Overall, the number of clusters obtained first increases, and then decreases with cutoff point, except in the high recombination rate scenario (Figure 3B). For each dataset we analyzed, the accuracy of clusters across similarity cutoffs was similar. However, the accuracy between different datasets varies, with higher average accuracy of clusters in datasets with high recombination rates (Figure 3C).

**Figure 3:**
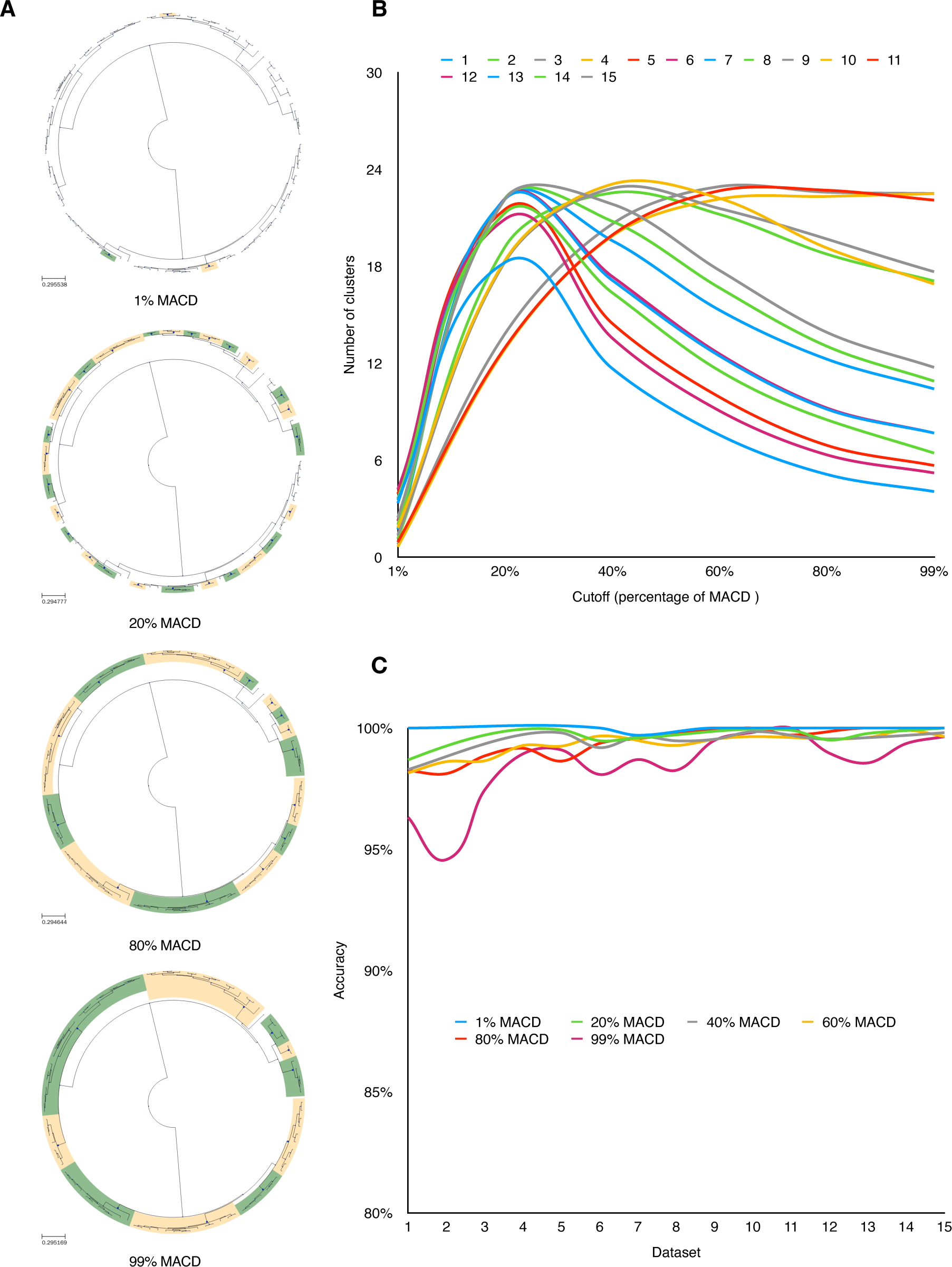
The performance of iterative-PopPUNK at multiple levels. A) the comparison of iterative-PopPUNK clusters with the genealogical tree (from 100 samples) at four levels: 1%, 20%, 80%, and 99% of the MACD. These green and orange faces represent matched clusters with tree nodes respectively. For these unmatched clusters, the tree leaves are annotated using different colors. B). The number of clusters iterative-PopPUNK tree for different cutoff values, averaged over datasets. C). The accuracy of estimated clusters at each level. The accuracy is calculated by the total number of matched clusters (with tree nodes) divided by the total number of estimated clusters. The chosen example dataset has a recombination rate r=0.006 and mutation rate μ=0.01

### Application of iterative-PopPUNK to real data shows consistency with phylogenetic results

We applied iterative-PopPUNK with default settings to analyze the genomic datasets of eight bacterial species with different mutation and recombination rates (Supplemental Table S1) and obtained the cluster assignments under six relative similarity cutoffs (P_max-cluster-dist_: 1%-99%). In addition, we performed core genome alignments for these datasets, and constructed maximum-likelihood (ML) phylogenetic trees based on core-genome SNPs. To measure the consistency between iterative-PopPUNK and alignment-based method, we mapped the clusters obtained under different cutoffs to the corresponding ML trees, and calculated matching accuracy (i.e. the proportion of matched clusters to all clusters) in the same manner as described above.

We collated the dataset for *Escherichia coli* by subsampling 500 genomes from 50 largest lineages (10 per lineage), which together represent the entire genomic diversity of human-isolated *E. coli* (Horesh et al. 2021). The entire, fully sampled, dataset has previously been analyzed by PopPUNK, and these 50 lineages were defined based on PopPUNK clusters. A maximum-likelihood (ML) phylogenetic tree based on core-genome SNPs was constructed for this same set of *E.coli* genomes to evaluate the matching rate of ST groups and this ML tree (Figure 4A).

**Figure 4:**
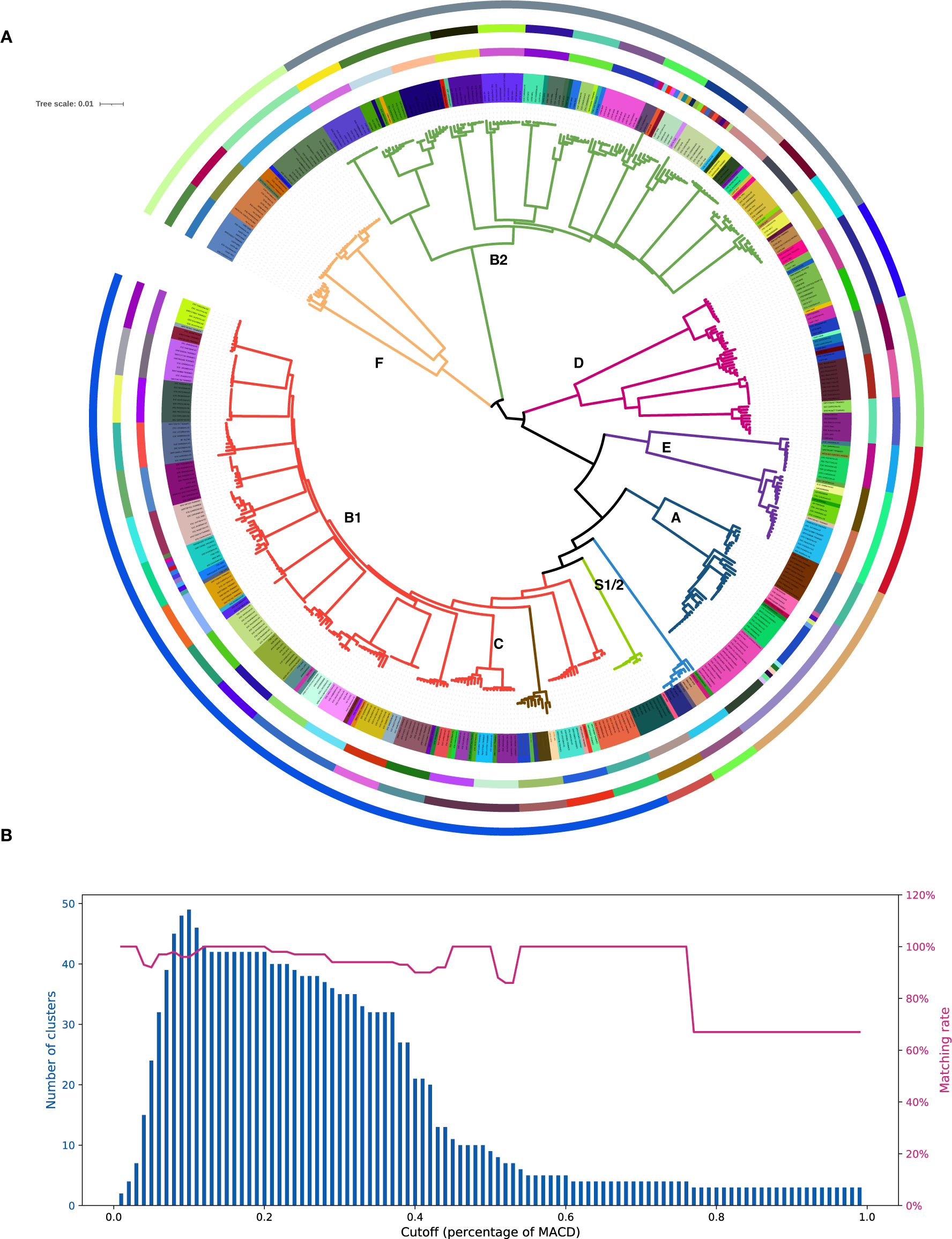
Population structure of E.coli at different iterative-PopPUNK levels. A). Comparison of PopPUNK clusters and a phylogenetic tree. The outer solid ring represents 9 PopPUNK clusters at 50% of the MACD, corresponding to 9 E.coli phylogroups. The middle ring contains 42 clusters at 20% of the MACD, corresponding to 42 clusters from phylogenetical tree at 20% SNP distance. The inner solid ring shows 94 PopPUNK clusters at 7.9% of the MACD, corresponding to 92 E.coli STs groups. Colors in the tree leaves and branches are used to present these 92 STs and 9 phylogroups respectively. The parameters used for estimating PopPUNK clusters were all set to default with BGMM model (two components). B). the distribution of matching rate and the number of clusters being estimated at multiple levels for this set of E.coli genomes. The tree was plotted using iTOL (https://itol.embl.de/). The comparison of PopPUNK clusters and an iterative-PopPUNK tree is available as Supplemental Fig S2. The original figure with high resolution is available as Supplemental Fig S3.

The result showed that these 55 STs groups from *E.coli* genomes were poorly coordinated with ML tree, as only 26 of them can exactly be matched to the tree nodes while the other 29 failed to find any corresponding nodes from the ML tree. The high mismatch rate between ST and ML tree is due to our very strict calculating method: these strains from unmatched clusters were closely related in the simulated tree, although not fully clustered together. However, there was nevertheless a high matching rates between ML tree and iterative PopPUNK clusters, with 75% being completely congruent (Figure 4B). If we look at the more permissive statistic of strain concordance (the total proportion of isolates that are assigned to one cluster by both methods), the matching rate is very high 91% (454/500) (Supplemental Fig S4).

Using a medium cutoff (P_max-cluster-dist_ =20%), iterative-PopPUNK estimated 42 clusters which exactly corresponded to 42 clusters using phylogenetic method with 20% SNPs distance as cutoff (Figure 4A). Under a large cutoff (P_max-cluster-dist_=50%), iterative-PopPUNK cluster closely corresponded to strains from a phylogroup. Of these 9 obtained clusters, Iterative-PopPUNK failed to find phylogroup C, which appears to be a sub-clade of B1 according to the phylogenetic tree and so does not deserve a phylogroup label according to current whole-genome analysis method. The definition of phylogroup C relied instead on virulence genes (Moissenet et al. 2010; Clermont et al. 2011; Clermont et al. 2013). Six perfectly corresponded to phylogroups (A, B2, E, F *Shigella flexneri and Shigella sonnei*) and the other two corresponded to the sub-clades of phylogroups D. This example highlighted the advantage of iterative-PopPUNK, which allows multi-level clustering in a unified way and users can select their “level” (e.g. sequence type or phylogroup level) according to their analytic purpose.

We also applied iterative-PopPUNK to analyse a “benchmark” dataset of *Vibrio parahaemolyticus*, consisting of 2,640 genomes, that had been previously analysed using phylogenomic methods (Yang et al. 2022). This dataset contains groups from clonal level (strains with ≤2500 SNPs) to outbreak level (within one-month clusters of strains with ≤6 SNPs) representing putative outbreaks. For this dataset, iterative-PopPUNK succeeded in estimating 23 clusters that were exactly corresponded to 23 pathogenic clonal groups (PCG) described in Yang et al. 2022 (Figure 5A; Supplemental Table S3). Two major PCGs, PCG3 and PCG189 (including 1,768 and 273 pathogenic genomes respectively) were extracted from this dataset to explore iterative-PopPUNK’s ability in resolving more fine-scale structure of closely related isolates. The results showed that iterative-PopPUNK clustering assignments had high concordance with outbreak-level phylogenomic clusters, with strains concordance of 90% (1371/1535) and 95% (259/273). However, as expected, the number of estimated “outbreak-level” iterative-PopPUNK clusters is less than that of phylogenomic clusters, because multiple phylogenomic clusters were assigned into one iterative-PopPUNK cluster. The number of clusters can be used as a rough quantification of the resolution. Using iterative PopPUNK, 80 clusters were identified for PCG3 and 25 clusters for PCG189. The concordance of iterative-PopPUNK clustering with the identified outbreak groups was 60% (48/80) for PCG3 and 80% (20/25) for PCG189. In both cases, the resolution is much finer than clonal level (Figure 5B and 5C; Supplemental Table S3).

**Figure 5.**
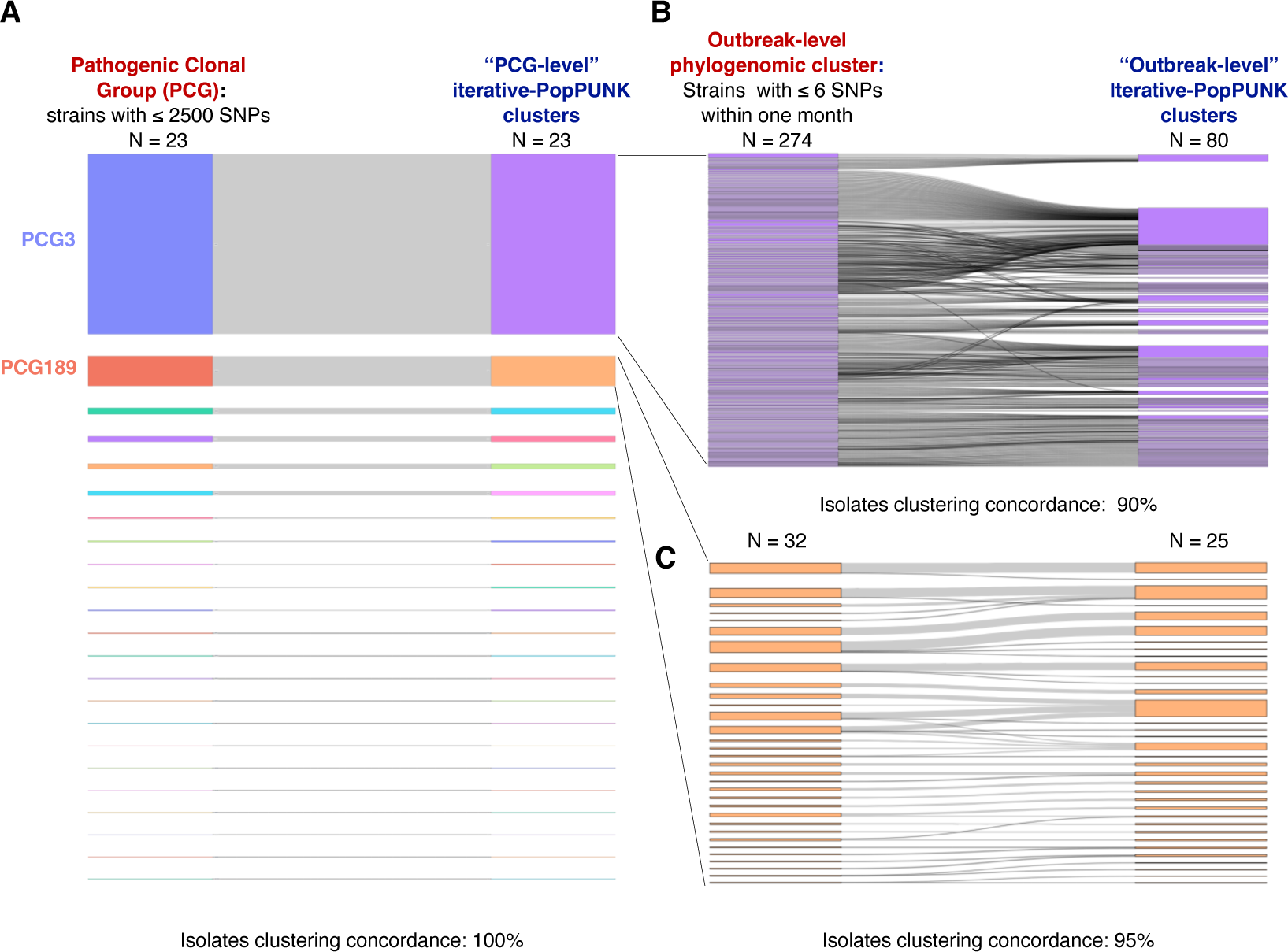
The comparison of Pathogenic Clonal Groups (PCG), outbreak-level phylogenomic clusters and iterative-PopPUNK cluster in Vibrio parahaemolyticus. (A) The comparison of PCGs described in Yang et al. 2022 and iterative-PopPUNK clusters. (B, C) The comparison of outbreak-level phylogenomic clusters of two major PCGs and iterative-PopPUNK clusters.

The datasets of the other six species were assembled by from publicly available genomes in NCBI RefSeq database. Iterative-PopPUNK did not produce useful results for two species with low genetic diversify, namely *Mycobacterium tuberculosis* and *Bacillus anthracis*. For the other four species, namely *Helicobacter pylori*, *Vibrio parahaemolyticus*, *Klebsiella pneumoniae* and *Bacillus cereus*, the average matching rates between phylogenetic methods and Iterative-PopPUNK were 85%, with higher rates in species with high recombination rates such as *Vibrio parahaemolyticus*. The matching rates did not show clear differences between at different similarity cutoffs (Figure 6, Supplemental Fig S5-S8).

**Figure 6:**
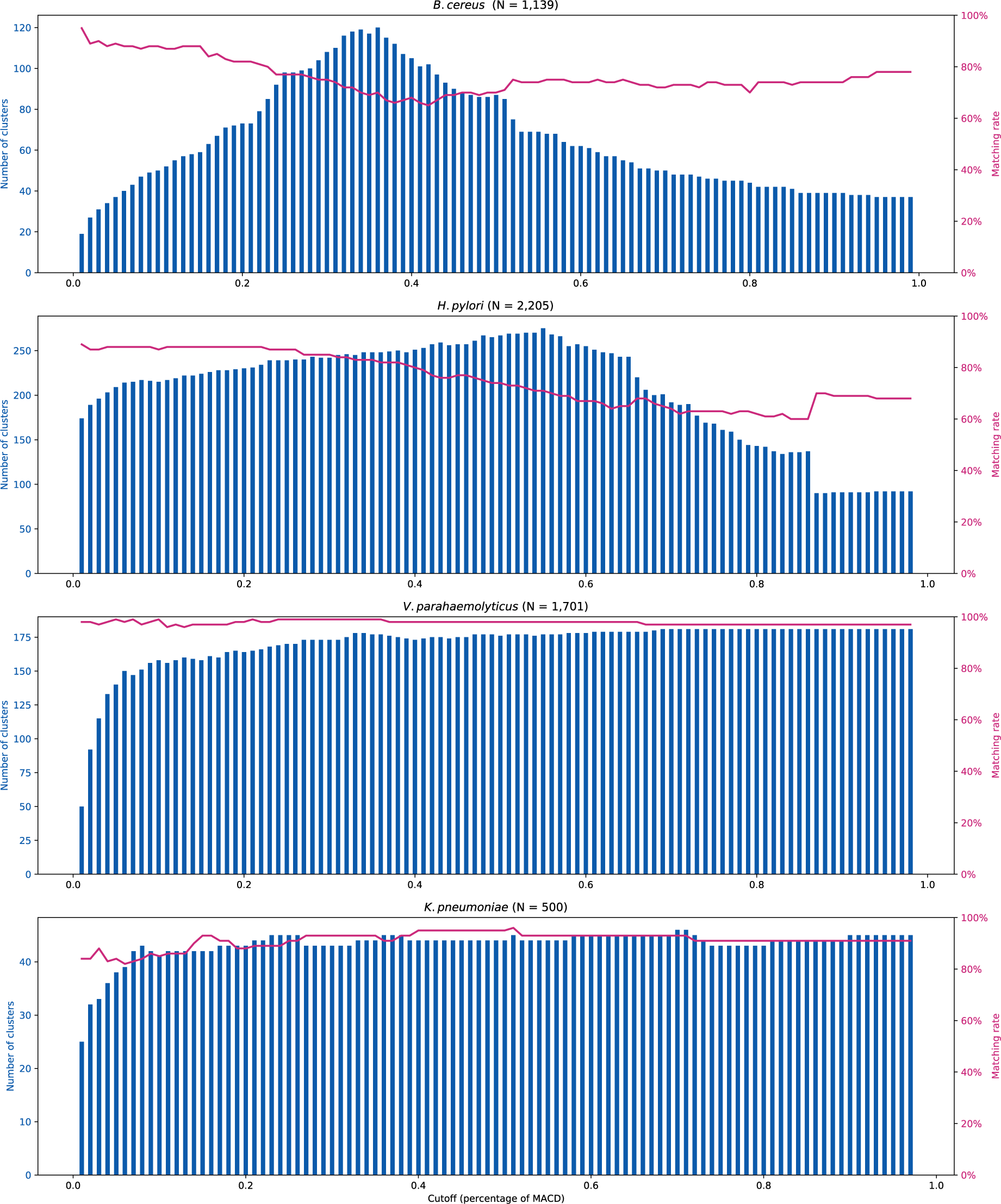
The distribution of matching rate and the number of clusters being estimated at multiple levels. A cluster is defined as a group with two or more strains (so non-clustered strains are not considered in counting the number of clusters)

## Discussion

Over the last two decades, genome data is accumulating at an unprecedented speed. For some common pathogens such as *Salmonella enterica* and *Escherichia coli*, there are now more than 200,000 genomes in public databases such as EnteroBase (Alikhan et al. 2018) and Genometracker (Brown et al. 2021). Analysing large-scale genome datasets is still challenging, and alignment-free methods such as PopPUNK are one of the best solutions to address this. PopPUNK was initially designed to identify a single level of clusters by optimising a heuristic, but in practice different cluster resolutions are required, depending on the purpose. To solve this problem, we developed iterative-PopPUNK, which allows estimation of a partially resolved genealogical tree and enables clustering at different levels of sequence similarity. Iterative-PopPUNK inherits the advantages of PopPUNK, which is fast and consumes few computational resources, and it is easily extendable when new data needs to be added (Lees et al. 2019). On this basis, Iterative-PopPUNK provides greater flexibility and allows the resolution of the clustering to be adjusted according to the specific purpose. However, default PopPUNK may be a better solution for analysing large datasets where a single level of clustering needs to be imposed, for example to define “non-redundant” genomes, that are representative of the species as a whole. In this case, the network score provides a mathematically founded method for choosing a “best” clustering, which is a pragmatic way of choosing a subset isolates that best represent the overall diversity of the dataset in a more compact form.

The traditional method for establishing the relationships between bacterial strains is phylogenetic analysis, where SNPs within a DNA sequence alignment are used to reconstruct a tree. These methods usually require a reference genome, and inappropriate reference selection may introduce bias. This approach is also not typically scalable to large-scale genome datasets (e.g. >10,000) and runtimes can become prohibitively long, even with generously provisioned computational resources. For example, on a high-end workstation a phylogenomic analysis of ∼10,000 *Klebsiella pneumoniae* genomes took 14 days and required a maximum of ∼500 GB memory. Furthermore, phylogenetic methods may be inaccurate in the presence of high homologous recombination rates, since individual recombination events can change multiple nucleotide positions within the alignment, making the accumulation of polymorphisms over time less clock-like. More sophisticated approaches that differentiate between recombination and mutation events can be even more time consuming.

Another kind of method for classifying strains is based on gene-based subtyping information, including the classically multi-locus sequence typing (MLST) which is based on several house-keeping genes (Maiden et al. 1998), and more recently developed core-genome MLST (cg-MLST) or whole-genome MLST (wg-MLST). Compared with genome-wide variation such as SNPs, subtyping information is typically much more parsimonious at cost of the loss of some phylogenetic resolution, making the analyses scalable to large datasets. HierCC is a state-of-the-art pipelines for multi-level clustering analysis of large-scale dataset based on cgMLST (Zhou et al. 2021). HierCC hierarchical clusters (HC) can be used for multi-level strain assignment. For example, in *Salmonella*, HC900 (<900 allele difference) and HC2500 (<2500 allele difference) represent strains from the same sequence type complex and species/subspecies, respectively. However, performing gene-by-gene subtyping-based clustering first requires an appropriate curated database of genes, and a stable strain nomenclature. These have not yet constructed for most bacterial species, especially for non-pathogenic species, and require ongoing maintenance, making this method unlikely to be the general solution for microbes. Both PopPUNK and Iterative-PopPUNK do not require a curated set of reference genomes and allow to train their own model based on any collection of genomes. The PopPUNK package does provide curated databases of several species, which users can use for quickly clustering new genomes.

The current version of iterative-PopPUNK has two main limitations. First, it does not perform well on species with low genetic diversity because boundaries tend to connect clusters together in a non-transitive manner. This can be solved by making a neighbor joining tree within each strain using the core distances calculated by PopPUNK. Second, the range of the levels of clusters obtained depends on the model fit of within- and between-strains components. Sometimes, only a very limited number of levels can be obtained, especially when using the second type of relative similarity cutoff P_bet-mean-dist_.

With the wide use of whole genome sequencing and rapid accumulation of genome sequences, large-scale genome dataset analysis may become normal in the near future. However, there is currently no ideal multi-level clustering method for large-scale dataset analysis. Iterative-PopPUNK is alignment-free, highly efficient even for very large-scale dataset, and does not rely on a curated set of reference genomes database and nomenclature, which can solve the limitations of the above methods, and we believe has can be a solution for fast multi-level clustering of most bacterial species. However, where data size and computation time is not an issue, methods that use alignments and evolutionary models to reconstruct the individual genetic events involved in strain divergence capture more information and have the potential to achieve slightly higher clustering resolution.

## Methods

### Implementation of iterative-PopPUNK algorithm

Iterative-PopPUNK procedure is implemented in the ‘poppunk_iterate’ program installed as part of PopPUNK package. Source code is available on GitHub (see Software availability) and can be installed through conda, pip or manually. Users can run multi-level clustering procedure using the flags ‘--fit-model refine--multi-boundary’ in the PopPUNK software.

The subsequent QC procedure is implemented in the ‘poppunk_iterate’ program and uses output from multi-level clustering step as input.

### Iterative-PopPUNK cluster accuracy evaluation using simulation data

We performed simulation using FastSimBac (De Maio and Wilson 2017) to test the accuracy of iterative-PopPUNK clusters. Fifteen parameter combinations were used in simulation to represent bacteria with different mutation and recombination rates (Supplemental Table S1). Simulation outputs were converted to genome sequences in FASTA format for further iterative-PopPUNK analysis using integrated simulator, *msformatter* and *ms2dna* (available from http://guanine.evolbio.mpg.de/bioBox), and the corresponding genealogical trees were extracted for comparison with iterative-PopPUNK clusters.

To analyze of simulated genomes, we set the maximum accessory distance parameter of Iterative-PopPunk to 1.0 to retain all isolates. In the first “model fitting” step, BGMM from original PopPUNK was chosen to fit the distribution of core and accessory distances. Two components (--K 2), one for within-strain and the other one for between-strain were applied. The component closest to the origin is defined as within-strain, in which all these shorter pairwise distances are from one cluster; while the between-strain distances refer to these larger ones that may be from different clusters. The other parameters were set to default values (k=15-29 in steps of 4; sketch size 10^4^), which have been shown to work well across most bacterial species.

Based on the node information of simulated genealogical trees, we assign simulated strains into true tree-clusters. Each iterative-PopPUNK cluster was compared with the true tree-clusters using ete3 toolkit (Huerta-Cepas et al. 2016). If the strain assignment of an iterative-PopPUNK cluster is identical to one of these simulated tree-clusters, it is defined as a match iterative-PopPUNK cluster, and associated nodes were labeled using green/red faces alternatively in the simulated tree; otherwise, it is unmatched, and the leaves would be annotated using a certain color for each unmatched cluster. Accuracy is defined as the proportion of matched clusters to all iterative-PopPUNK clusters. The method of Robinson and Folds (Robinson and Foulds 1981) were applied to compare the simulated genealogical tree and the inferred iterative-PopPUNK tree using the ete3 toolkit in Python.

### Application of iterative-PopPUNK in real data

The publicly available, assembled genomes of seven species were downloaded from the National Center for Biotechnology Information (NCBI) Reference sequence (RefSeq) database based on the taxonomy identifier (Supplemental Table S2). Two steps of genome sequences QC were performed. First, we used CheckM (Parks et al. 2015) to assess the genome quality and only genome sequences with completeness >90% and contamination <5% were kept in further analysis. Second, we set 70% as whole genome coverage threshold to remove low-quality genomes or outlier samples that may not be part of the species. After QC, the same iterative-PopPUNK pipeline used for simulation data were applied to real data analysis. The 500 *E. coli* genomes were collected from Horesh G et al. Microb Genom. 2021. Additionally, a separate set of benchmark data comprising 2,640 *Vibrio parahaemolyticus* genomes (distinct from those obtained from the NCBI database) was obtained from Yang et al. Nat Microb. 2022. The original dataset consisted of 3,460 genomes, but singletons and non-pathogenic isolates were removed for this analysis. The Sankey Diagrams were plotted using Plotly graphing library for Python (available from https://github.com/plotly/plotly.py).

QC-passed genome sequences were further aligned to the reference genome of each species (Supplemental Table S2) using MUMmer (Kurtz et al. 2004) to generate whole genome alignments. Core-genome (regions present in >99% isolates) SNPs were called using snp-sites (Page et al. 2016) based on the alignments. ML phylogenetic tree were constructed using FastTree (Price et al. 2010) based on core-genome SNPs. We adopted the method used in Iterative-PopPUNK cluster accuracy evaluation to calculate the matching rate of iterative-PopPUNK clusters and ML trees, by replacing the simulated trees to ML trees. In order to show how the number of estimated clusters changes for different species, we used 1% MACD as a gap to move the cutoff from 1% MACD to 99% MACD. For practical usage of iterative-PopPUNK, users can adjust these cutoffs as needed.

### Software availability

The source code and tutorial are available in Supplemental Code. The program ‘poppunk_iterate.py’ is available on GitHub (https://github.com/bacpop/PopPUNK), installed as a part of PopPUNK package. Documentation for tutorial can also be found on https://poppunk.readthedocs.io/en/latest/poppunk_iterate.html. Online examples from tutorial for each dataset are listed in Supplemental Table S3

## Competing interest statement

The authors declare no competing interests.

## Acknowledgments

This work is supported by the National Key Research and Development Program of China (No. 2022YFC2304700), Shanghai Municipal Science and Technology Major Project (No. 2019SHZDZX02) to DF. CY is funded by National Natural Science Foundation of China (No. 32270003, No. 32000008), Youth Innovation Promotion Association, Chinese Academy of Sciences (No. 2022278) and Shanghai Rising-Star Program (No. 23QA1410500). JAL received support from the Medical Research Council (grant number MR/R015600/1). This award is jointly funded by the UK Medical Research Council (MRC) and the UK Department for International Development (DFID) under the MRC/DFID Concordat agreement and is also part of the EDCTP2 program supported by the European Union.

## Notes

### Competing Interest Statement

The authors have declared no competing interest.

### Summary of Updates

The revised manuscript includes several important updates and improvements. Firstly, in the introduction section, we introduced the algorithm of PopPUNK, which can be found on Page 4, Lines 75-81 of the revised manuscript. Secondly, Figure 1 and its legend have been further expanded to enhance clarity and improve understanding. Thirdly, we added the Average Robinson Folds distance between true trees and iterative-PopPUNK trees to Figure 2E, providing valuable results for analysis. In terms of documentation, the iterative-PopPUNK tutorial section has been substantially improved, offering a detailed explanation of the approach. Additionally, several example datasets have been included to assist users in understanding and utilizing the method. The latest version of the iterative-PopPUNK tutorial can be accessed at https://poppunk.readthedocs.io/en/latest/poppunk_iterate.html. Furthermore, comparisons between iterative-PopPUNK and phylogenetic trees for real data have been included in Supplemental Figures S5-S8, providing further insights. To ensure accuracy, the revised manuscript has been carefully checked by a native speaker, who has corrected any typographical errors. Additionally, iterative-PopPunk has been applied to a "benchmark" outbreak dataset of Vibrio parahaemolyticus, previously analyzed using phylogenomic methods. Figure 1C has undergone revision to provide a clearer representation of the percentage of MACD. Lastly, Figure 4B has been moved to Supplemental Figure S2, while Figure 4A has been zoomed in to enhance focus. Detailed information can also be found in Supplemental Table S3.

