## Supplemental Figure S1 for "Genealogical inference and more flexible sequence clustering using iterative PopPUNK"

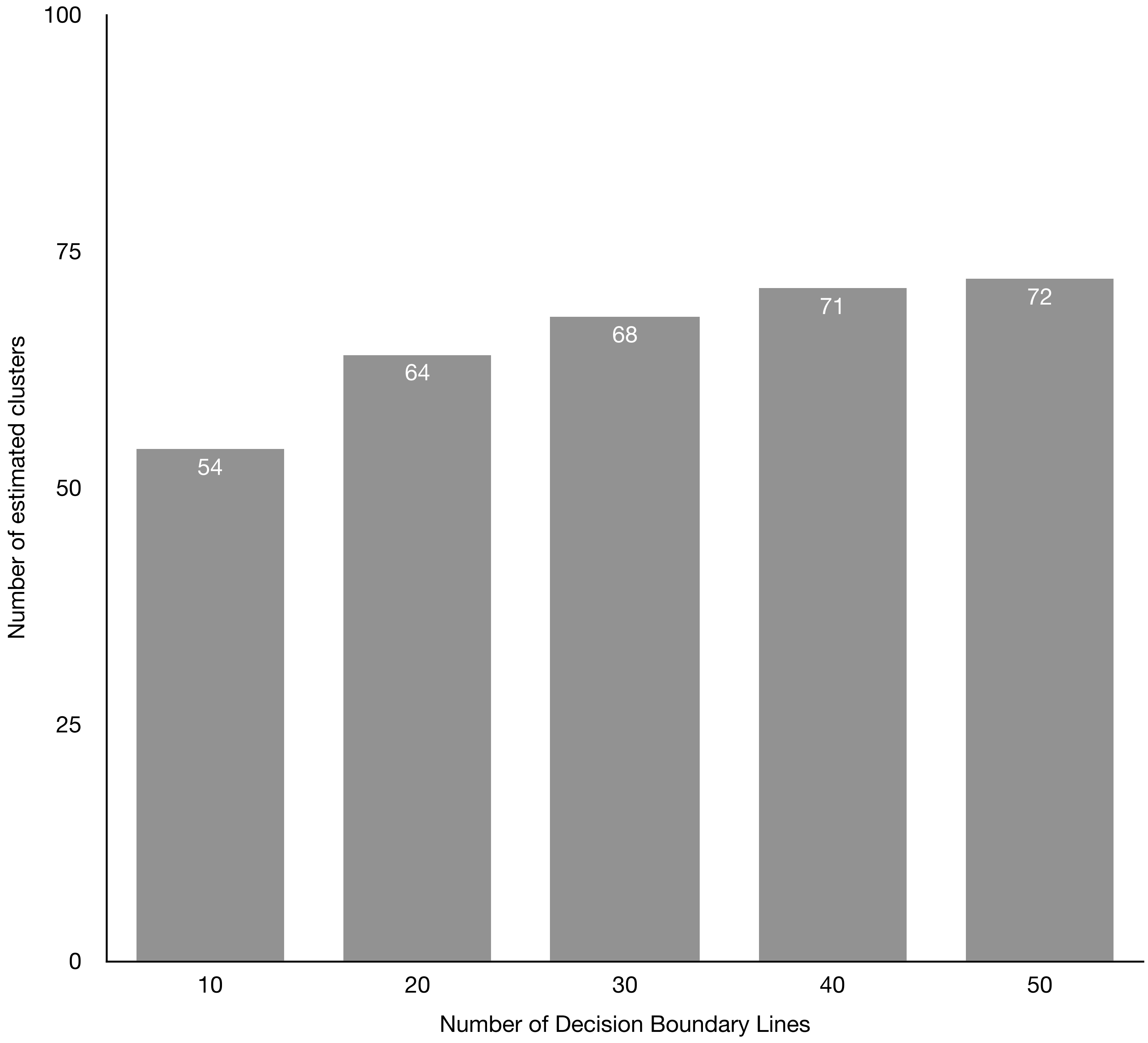


Supplemental Fig S1: The relationship between the number of decision boundary lines and estimated clusters. In this plot, the average number of estimated clusters were generated from 150 randomly selected sets of simulation genomes.
