## Supplemental Figure S3 for "Genealogical inference and more flexible sequence clustering using iterative PopPUNK"

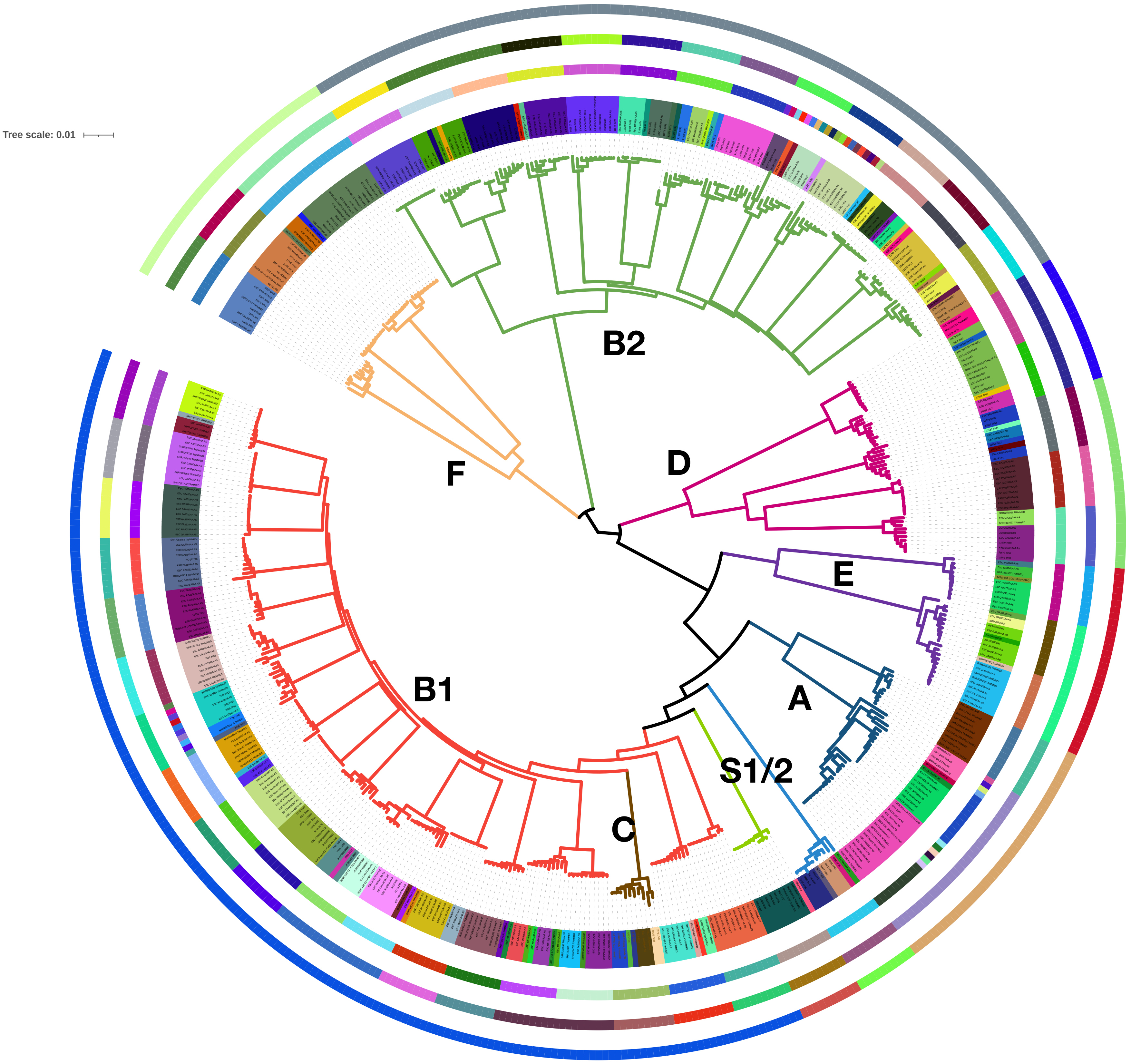


Supplemental Fig S3: Comparison of phylogenetic tree and PopPUNK clusters. The outer ring represents 9 PopPUNK clusters at 50% of the MACD, corresponding to 9 *E.coli* The middle ring contains 42 clusters at 20% of the MACD, corresponding to 42 clusters from phylogenetical tree at 20% SNP distance. The inner solid ring shows 94 PopPUNK clusters at 7.9% of the MACD, corresponding to 92 *E.coli* STs groups. Colors in the tree leaves and branches are used to present these 92 STs and 9 phylogroups respectively. The parameters used for estimating PopPUNK clusters were all set to default with 2D GMM model (two components) with a “neg-shift” value of -0.25.
