## Supplemental Figure S4 for "Genealogical inference and more flexible sequence clustering using iterative PopPUNK"

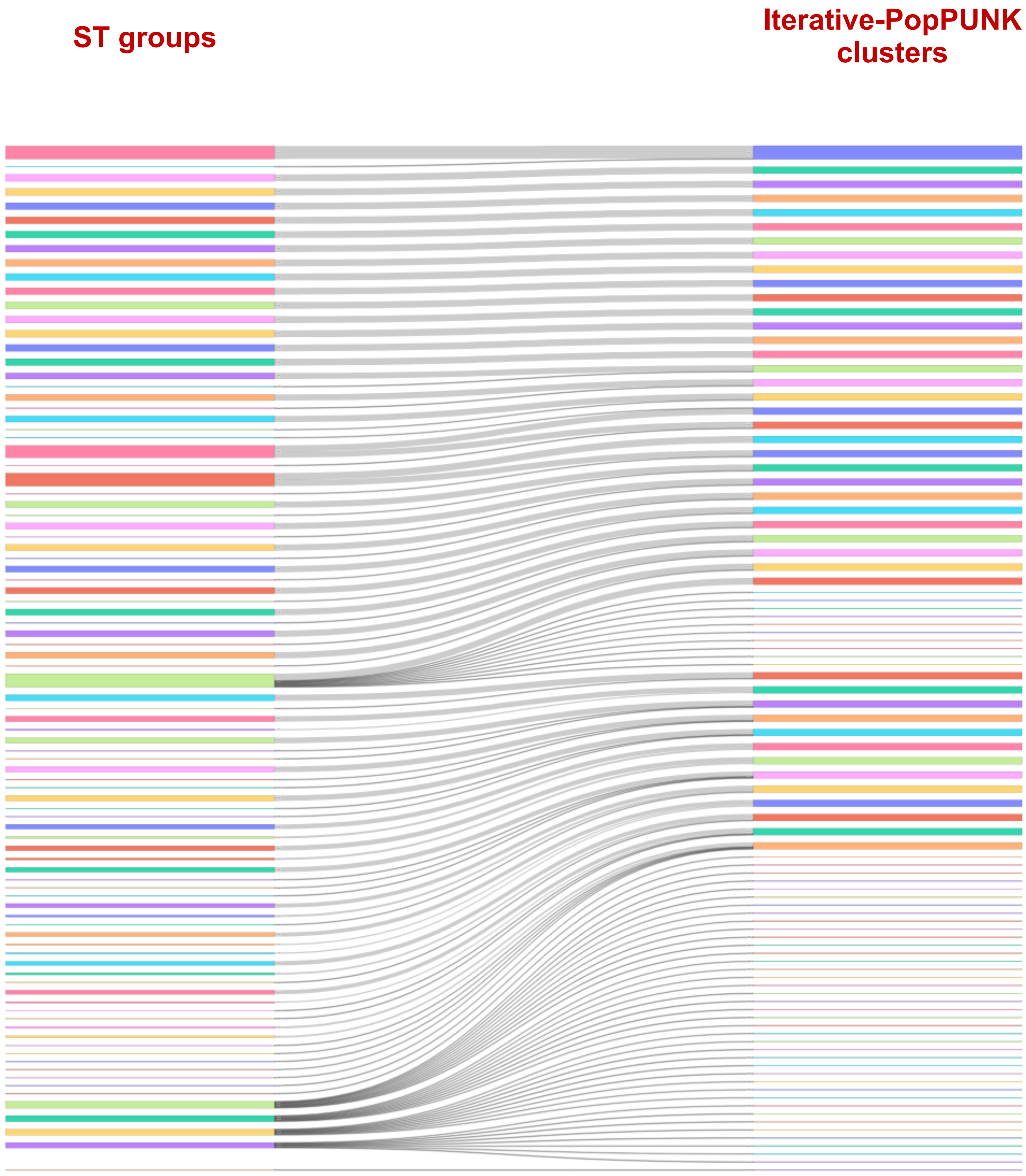


Supplemental Fig S4: The comparison of Sequence Typing (ST) groups and iterative-PopPUNK clusters of 500 representative *E.coli* genomes. The ST groups of these 500 *E.coli* genomes were extracted from Horesh et al. 2021 and the corresponding iterative-PopPUNK cutoff value is 7.9% of MACD.
