## Supplemental Figure S7 for "Genealogical inference and more flexible sequence clustering using iterative PopPUNK"

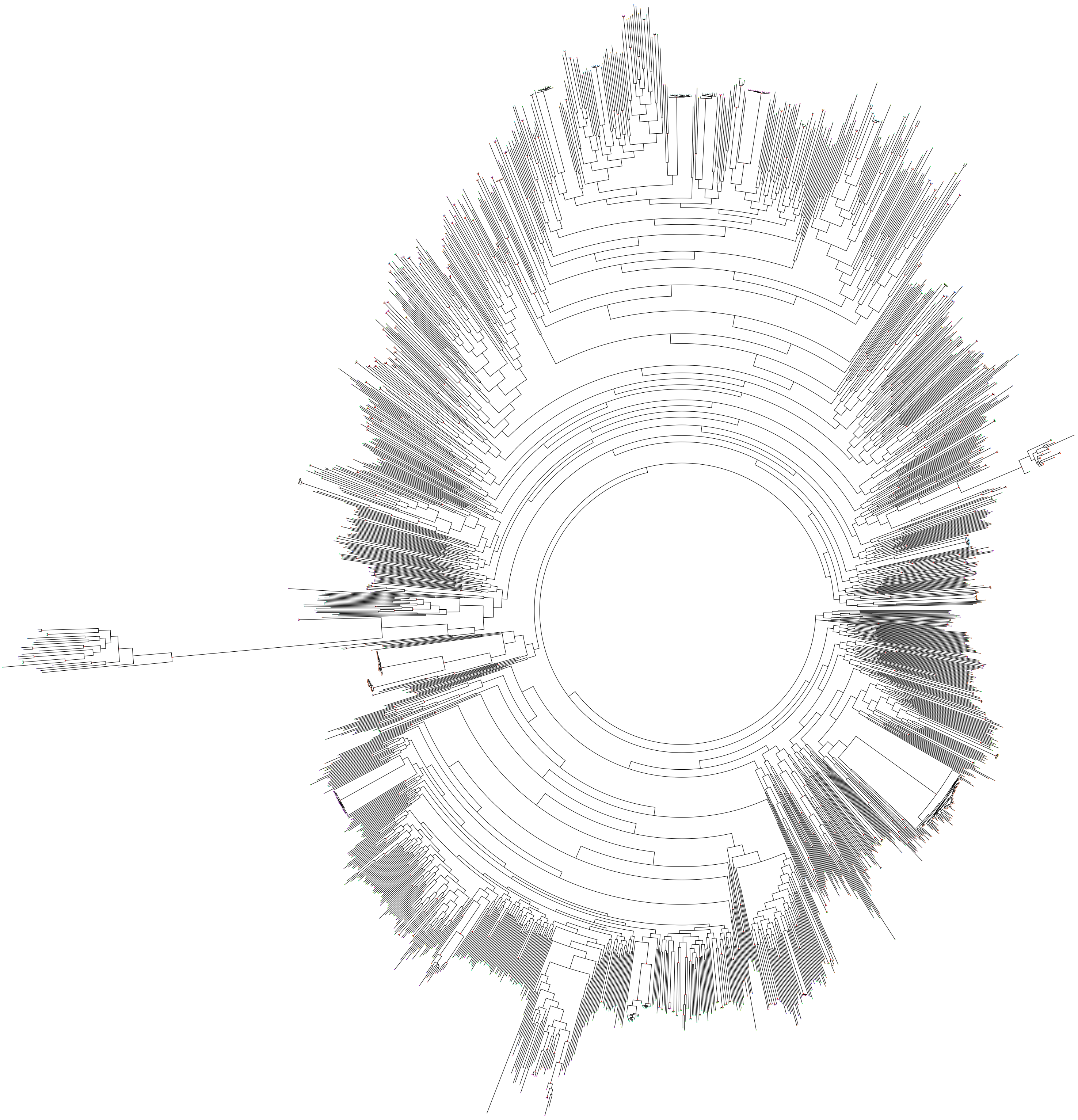

Supplemental Fig S7: The iterative-PopPUNK tree overlaid with and phylogenetic tree of *Helicobater pylori*. Nodes of the tree that correspond exactly to an inferred cluster are shown in red. Where an inferred cluster does not correspond to a node in the phylogenetic tree, it is assigned a colour and all the strains in that cluster have their tips labelled using that colour.
