## Supplemental Table S1 for "Genealogical inference and more flexible sequence clustering using iterative PopPUNK"

*Table S1: Parameters used in FastSimBac program to generate simulated genomes. Under each condition, A total of 100 simulated genomes were produced with 100 sequences in each genome. True clonal genealogies were also produced for each genome.*

| **Dataset** | **Total simulated genomes** | **Number of generated genomes per simulation** | **Sequence length** | **Recombination rate** | **Mutation rate** | **R/m** |
| --- | --- | --- | --- | --- | --- | --- |
| 1 | 100 | 100 | 1,000,000 | 0.003 | 0.01 | 0.30 |
| 2 | 100 | 100 | 1,000,000 | 0.006 | 0.01 | 0.60 |
| 3 | 100 | 100 | 1,000,000 | 0.01 | 0.01 | 1.00 |
| 4 | 100 | 100 | 1,000,000 | 0.02 | 0.01 | 2.00 |
| 5 | 100 | 100 | 1,000,000 | 0.04 | 0.01 | 4.00 |
| 6 | 100 | 100 | 1,000,000 | 0.003 | 0.03 | 0.10 |
| 7 | 100 | 100 | 1,000,000 | 0.006 | 0.03 | 0.20 |
| 8 | 100 | 100 | 1,000,000 | 0.01 | 0.03 | 0.33 |
| 9 | 100 | 100 | 1,000,000 | 0.02 | 0.03 | 0.67 |
| 10 | 100 | 100 | 1,000,000 | 0.04 | 0.03 | 1.33 |
| 11 | 100 | 100 | 1,000,000 | 0.003 | 0.06 | 0.05 |
| 12 | 100 | 100 | 1,000,000 | 0.006 | 0.06 | 0.10 |
| 13 | 100 | 100 | 1,000,000 | 0.01 | 0.06 | 0.17 |
| 14 | 100 | 100 | 1,000,000 | 0.02 | 0.06 | 0.33 |
| 15 | 100 | 100 | 1,000,000 | 0.04 | 0.06 | 0.67 |
