## Supplemental Table S2 for "Genealogical inference and more flexible sequence clustering using iterative PopPUNK"

*Table S2: Summary of real data*

| **Species** | **Taxonomy ID** | **Total genome sequences** | **Genome sequences (after checkm)** | **Sequences with coverage problem** | **Sequences (after QC)** |
| --- | --- | --- | --- | --- | --- |
| *Vibrio parahaemolyticus* | 670 | 1,748 | 1,702 | 1 | 1,701 |
| *Helicobacter pylori* | 210 | 2,241 | 2,205 | 0 | 2,205 |
| *Bacillus cereus* | 1396 | 1,220 | 1,205 | 66 | 1,139 |
| *Bacillus anthracis* | 1392 | 336 | 333 | 0 | 333 |
| *Klebsiella pneumoniae* | 573 | 500 | 500 | 0 | 500 |
| *Mycobacterium tuberculosis* | 1773 | 6,951 | 6,934 | 0 | 6,934 |
